# Predictive and Interpretable: Combining Artificial Neural Networks and Classic Cognitive Models to Understand Human Learning and Decision Making

**DOI:** 10.1101/2023.05.17.541226

**Authors:** Maria K. Eckstein, Christopher Summerfield, Nathaniel D. Daw, Kevin J. Miller

## Abstract

Quantitative models of behavior are a fundamental tool in cognitive science. Typically, models are hand-crafted to implement specific cognitive mechanisms. Such “classic” models are interpretable by design, but may provide poor fit to experimental data. Artificial neural networks (ANNs), on the contrary, can fit arbitrary datasets at the cost of opaque mechanisms. Here, we adopt a hybrid approach, combining the predictive power of ANNs with the interpretability of classic models. We apply this approach to Reinforcement Learning (RL), beginning with classic RL models and replacing their components one-by-one with ANNs. We find that hybrid models can provide similar fit to fully-general ANNs, while retaining the interpretability of classic cognitive models: They reveal reward-based learning mechanisms in humans that are strikingly similar to classic RL. They also reveal mechanisms not contained in classic models, including separate reward-blind mechanisms, and the specific memory contents relevant to reward-based and reward-blind mechanisms.

## Introduction and Background

Computational models of behavior are a fundamental tool in many areas of cognitive science. Some of the most popular models include reinforcement learning (RL; Daw, 2011; R. R. Miller et al., 1995; Sutton and Barto, 2017), Bayesian inference (Ma, 2019), and evidence accumulation (Ratcliff and McKoon, 2008). Computational models have a long and rich history in the cognitive sciences, partly because they hold the tantalizing promise to quantitatively test elaborate hypotheses about unobservable cognitive processes.

To this aim, computational models typically embody handcrafted hypotheses about a cognitive process of interest (Busemeyer and Diederich, 2010). For example, RL models assume that long-term reward histories are compressed into a low-dimensional, Markovian value representation, which is incrementally updated after new reward experiences, following a delta rule (Daw, 2011; Sutton and Barto, 2017). However, such classical models face problems (Eckstein et al., 2021):

1. It is fundamentally unclear *how well* any given model fits the data (Box, 1979; Navarro, 2019). Using the classic approach, several competing models are usually compared in terms of fit to identify the best (Katahira, 2016; Wilson and Collins, 2019). However, it is unclear when to stop searching: How do we determine that a winning model fits the data “well enough” (Palminteri et al., 2017)?
2. There also is the danger of model misspecification, e.g., of overlooking a mechanism that is crucial in the true data generating process (Nassar and Frank, 2016; Nussenbaum and Hartley, 2019). If a model is misspecified, we are not only disregarding a crucial mechanism, but other, correctly specified mechanisms will compensate for the deficit, leading to additional distortions (Sugawara and Katahira, 2021).

In this study, we argue that deep learning, amongst many possible applications in the cognitive sciences (Barak, 2017; Ma and Peters, 2020), can address these issues: 1) The universal function approximation theorem states that any sufficiently powerful artificial neural network (ANN) can approximate any input-output function arbitrarily well (Cybenko, 1989; Hornik, 1991): Sufficiently deep ANNs, trained on sufficiently large datasets, should therefore be able to represent any computational processes (LeCun et al., 2015), including cognitive computational processes. Recent studies have put this claim to test, and indeed found near-optimal fits of ANNs in simulation (Ger et al., 2023). In empirical data, many studies have shown extensive improvements in fit compared to hand-crafted models (e.g., Agrawal et al., 2020; Battleday et al., 2020; Kuperwajs et al., 2022; Peterson et al., 2021; Sutskever and Nair, 2008), including in RL tasks (Dezfouli, Ashtiani, et al., 2019; Dezfouli, Griffiths, et al., 2019; Fintz et al., 2021; Jaffe et al., 2022; Song et al., 2017; Yang et al., 2019).

2) ANNs offer the promise of overcoming model misspecification, precisely because they do not put constraints on the underlying mechanism, and hence allow any mechanism to be discovered. However, this flexibility also is a drawback: ANNs are often called “black boxes” because they do not— unlike hand-crafted models—offer direct insight into the process they learn to emulate (Ma and Peters, 2020).

Here, we first fit a classic hand-crafted RL model and an ANN to the same human learning task (Bahrami and Navajas, 2020). We formulate the RL model as a special case of the general ANN, which allows us to construct “hybrid” RL-ANN models to fill the gaps between both extremes: We will endow the RL model—step-by-step—with specific, interpretable, and delineated mechanisms until we obtain the fully general ANN. In this process, we calculate model fits to assess the empirical evidence for each added mechanism in human behavior. We also probe and inspect the component mechanisms of each fitted model: For example, we can directly compare an ANN’s learned input-output mappings to the fixed input-output mappings of the classic RL model. The main contribution of this study is to create cognitive models that capture human learning exhaustively, yet provide insight into its mechanisms.

## Results

### Dataset

#### Human Task

In a publicly available dataset, 965 human participants completed 150 trials of a drifting 4-armed bandit task (Bahrami and Navajas, 2020; original task by Daw et al., 2006). On each trial, participants were asked to select one of four bandits and received a continuous numerical reward (1-98 points) based on the chosen bandit’s current reward payout. Reward payouts drifted over time (Gaussian walk). Each participant was randomly assigned to one of three predetermined reward payoff schedules. For details, refer to Bahrami and Navajas (2020) and Daw et al. (2006).

#### Previous Findings

The task was originally designed to study the neural mechanisms underlying exploratory and exploitative choice (Daw et al., 2006). More recently, Fintz et al. (2021) trained ANN models that suggest that participants used RL-like, reward-sensitive as well as exploration-like, reward-insensitive strategies. However, the study did not address how both mechanisms interact, a question we will address here.

#### Data Preprocessing

We removed all participants who had missed more than 10% of trials (57 participants, 5.9%), a visually-determined elbow point, for a final sample size of 908. We set aside 20 participants per payoff schedule (60 total) as a testing dataset to assess model fit. Of the remaining 848 participants, we selected the largest balanced sub-sample in terms of reward schedules (825 participants; 275 per schedule) for the training dataset. Reward magnitudes were divided by 100 (resulting range: 0.01-0.98).

### Modeling

#### Modeling Strategy

In its basic form, the classic RL model RL_αβ_ comprises two functions: A classic, fixed value update rule *f*_*RL*_ that calculates action values *v*(*a*) based on observed rewards *r*_*t*_; and a softmax action rule that translates values *v*(*a*) into action probabilities *p*(*a*) (Table 1). Our first hybrid model replaces the classic, fixed value update *f*_*RL*_ with a recurrent neural network (RNN; a sequential ANN) *f*_*RNN*_. This allows the shape of the human learning rule to be learned empirically. Our next models expand the list of inputs to the value update *f*_*RNN*_ to assess which information humans use to calculate values *v*(*a*). For example, are value updates contextualized by the current value landscape, or is the previous value of an action enough to determine its new value? Lastly, we test if other processes besides reward-based learning affect human choice. We construct an ANN that calculates reward-independent “choice kernels”, and explore the interactions between value and choice kernel.

**Table 1:** Specification of all models in terms of their update and action rules. “Model Names” (left column) are structured as follows: “RL_αβ_” refers to the basic, classic RL model. Model names containing “RNN” [“LSTM”] refer to models that replace the classic RL update rule with a flexible RNN [LSTM]: Their update rules (right column) use *f*_*RNN*_ [*f*_*LSTM*_] rather than *f*_*RL*_. “O-”, “S-”, and “OS-” (left column) indicate the memory-based inputs to each model’s update rule (right column): “O-” and “OS-” models have access to their own previous output [*v, c*, or *g*]; “S-” and “OS-” models have access to their previous hidden state *s*. “-v” and “-c” (left column) indicate the observation-based inputs to each model’s update rule (right column): rewards *r* allow for the calculation of values *v*; and choices *a* for choice-kernels *c*. “-cv” refers to models that track both *v* and *c*, and combine them at decision time: The action rule (right column, top row in each panel) in “v” [“c”] models is based on just values *v* [choice kernels *c*]; “cv” models calculate both and combine them additively (i.e., without the capability to show *c*-*v* interactions). “-all” indicates models that have access to all available information simultaneously, allowing them to model *c*-*v* interactions. Best-fitting models are in bold.

| Model Names | Update and Action rules |
| --- | --- |
| *Value (“v”) Models* | $p_t(a) = \text{softmax}(v_t(a))$ |
| $RL_{\alpha\beta}$ | $v_{t+1}(a_t) += f_{RL}(v_t(a_t), r_t)$ |
| RNN-v | $v_{t+1}(a_t) += f_{RNN}(v_t(a_t), r_t)$ |
| O-RNN-v | $v_{t+1}(a_t) += f_{RNN}(v_t(a_t), r_t, v_t)$ |
| S-RNN-v | $v_{t+1}(a_t) += f_{RNN}(v_t(a_t), r_t, s_t)$ |
| OS-RNN-v | $v_{t+1}(a_t) += f_{RNN}(v_t(a_t), r_t, v_t, s_t)$ |
| LSTM-v | $v_{t+1}(a_t) += f_{LSTM}(v_t(a_t), r_t, s_t)$ |
| *Choice-Kernel (“c”) Models* | $p_t(a) = \text{softmax}(c_t(a))$ |
| RNN-c | $c_{t+1}(a) = f_{RNN}(a_t)$ |
| O-RNN-c | $c_{t+1}(a) = f_{RNN}(a_t, c_t)$ |
| S-RNN-c | $c_{t+1}(a) = f_{RNN}(a_t, s_t)$ |
| OS-RNN-c | $c_{t+1}(a) = f_{RNN}(a_t, c_t, s_t)$ |
| LSTM-c | $c_{t+1}(a) = f_{LSTM}(a_t, c_t, s_t)$ |
| *Additive (“cv”) Models* | $p_t(a) = \text{softmax}(v_t(a) + c_t(a))$ |
| RNN-cv | $v_{t+1}(a_t) += f_{RNN}(v_t(a_t), r_t)$<br>$c_{t+1}(a) = f_{RNN}(a_t)$ |
| O-RNN-cv | $v_{t+1}(a_t) += f_{RNN}(v_t(a_t), r_t, v_t)$<br>$c_{t+1}(a) = f_{RNN}(a_t, c_t)$ |
| S-RNN-cv | $v_{t+1}(a_t) += f_{RNN}(v_t(a_t), r_t, s_t)$<br>$c_{t+1}(a) = f_{RNN}(a_t, s_t)$ |
| **OS-RNN-cv** | $v_{t+1}(a_t) += f_{RNN}(v_t(a_t), r_t, v_t, s_t)$<br>$c_{t+1}(a) = f_{RNN}(a_t, c_t, s_t)$ |
| **LSTM-cv** | $v_{t+1}(a_t) += f_{LSTM}(v_t(a_t), r_t, s_t)$<br>$c_{t+1}(a) = f_{LSTM}(a_t, c_t, s_t)$ |
| *Interactive (“all”) Models* | $p_t(a) = \text{softmax}(g_t(a))$ |
| $RL_{\alpha\beta pf}$ | $g_{t+1}(a_t) += f_{RL}(v_t(a_t), r_t, a_t)$ |
| RNN-all | $g_{t+1} = f_{RNN}(r_t, a_t)$ |
| O-RNN-all | $g_{t+1} = f_{RNN}(r_t, a_t, g_t)$ |
| S-RNN-all | $g_{t+1} = f_{RNN}(r_t, a_t, s_t)$ |
| OS-RNN-all | $g_{t+1} = f_{RNN}(r_t, a_t, g_t, s_t)$ |
| **LSTM-all** | $g_{t+1} = f_{LSTM}(r_t, a_t, s_t)$ |

#### Classic Cognitive Models: “RL_αβ_” and “RL_αβ*p f*_”

RL models are built on the notion of values and reward prediction errors (Daw, 2011; Wilson and Collins, 2019): Learners acquire, through trial-and-error, a value *v*(*a*_*i*_) for each action *a*_*i*_. Values are acquired incrementally, by updating them each time a reward *r* is observed, based on the reward prediction error *rpe* = *r* −*v*(*a*_*i*_). To update *v*(*a*_*t*_), having observed *r*_*t*_, previous values are moved slightly in the direction of the observed outcome: *v*_*t*+1_(*a*_*t*_) = *v*_*t*_(*a*_*t*_) +α\**rpe*, with the learning rate parameter α controling the update size. To select actions based on these (potentially continuously evolving) values, the cognitive literature often employs the “softmax” transform: *p*(*a*) = *so f tmax*(β* *v*(*a*)).^1^ The inverse decision temperature parameter β controls the stochasticity.

To fit our RL_αβ_ model to the dataset, we initialized action values at 0.5 and performed gradient descent (Adam optimizer) on the cross-entropy loss 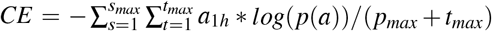. *a*_1*h*_ is the one-hot vector of participants’ actions; *p*(*a*) are model-derived action probabilities for each subject *s* and trial *t* (batch size *s*_*max*_ = 64). Parameters α = 0.77 and β = 7.05 led to the best fit on the training data. (For testing loss and prediction accuracy on held-out participants, see Table 1 and Fig. 1B).

**Figure 1:**
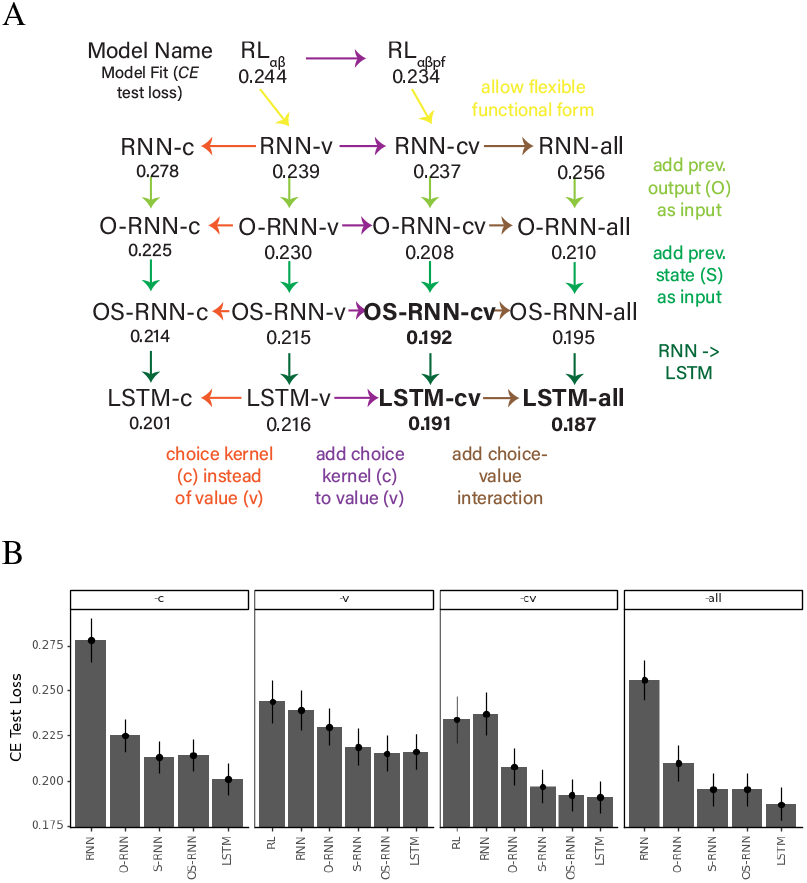
Overview and fit of the main models. A) Models are arranged according to their complexity, starting with *RL*_αβ_ at the top-left, and ending with LSTM-all at the bottom-right. Three main features differentiate *RL*_αβ_ from LSTM-all: The ability to flexibly learn the update rule from data (yellow, diagonal); additional memory capacity (green hues, vertical); and the existence of a reward-blind choice mechanism, and its interaction with the reward-based mechanism (red hues, horizontal). Arrow colors show which mechanism is added for each model. Numbers underneath model names indicate model fit (explained in panel B). Best-fitting models in bold. B) Model fits, i.e., *CE* losses on the held-out test dataset (lower is better). Each panel shows one model family (“-c”: Choice-kernel models; “-v”: value models; “-cv”: additive models; “-all”: interactive models; see Table 1).

To assess qualitative model fit (Palminteri et al., 2017), we characterized models’ ability to reproduce human behavior: We simulated behavior from the best (in training) among four models, asking the model to select actions based on its own proposed action probabilities, using human-fitted parameters. We simulated *n* = 825 agents with *t* = 150 trials, and subjected the simulated agents to the same behavioral analyses as humans. Although the RL_αβ_ model learned to perform the task, it did not reproduce typical human behavioral patterns (Fig. 2B). To address this shortcoming, the RL_αβ_ model is commonly expanded, for example by forgetting and choice persistence mechanisms, parameterized by *f* and *p*, respectively (e.g., Eckstein et al., 2022). Adding these parameters in model RL_αβ*p f*_ led to a slight improvement in model fit (Fig. 1).

**Figure 2:**
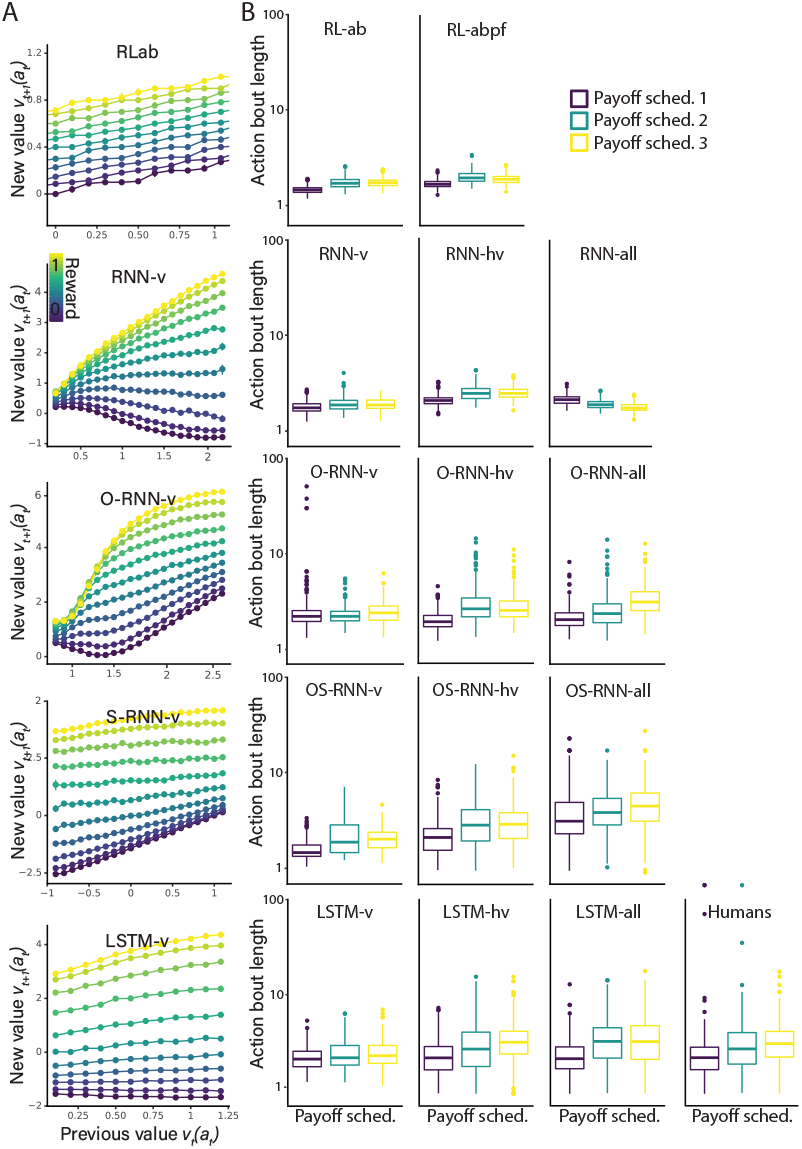
A) Analyzing learned learning rules. All “v models” update *v*(*a*) on each trial, using parameters and (except for RL_αβ_) update functions (learning rules) learned from human data. X-axes (*v*_*t*_(*a*_*t*_)) and colors (*r*_*t*_) show function inputs, y-axes show function outputs (*v*_*t*+1_(*a*_*t*_)), for a full specification of the learned functions. Methods: 10.000 random input tuples were passed through each update function to obtain outputs. *r*_*t*_ was sampled randomly across the allowed range [0, 1]; *v*_*t*_(*a*_*t*_) and *s*_*t*_ were sampled within 3 standard deviations of their mean along their first principal component (PC). Points indicate means, error bars standard errors of the mean, after binning data points into 0.1-sized bins based on reward magnitude. Differences in y-axis scale can be interpreted as differences in “decision temperature”. B) Assessing qualitative model fit using behavioral simulations. We used human-fitted parameters θ to simulate artificial behavior, and subjected all (artificial and human) datasets to the same behavioral analyses. Action bout length (participants’ average numbers of subsequent identical actions) is one example behavior. Box plots show the 25th, 50th (median), and 75th percentile over participants; whiskers extend to the 1.5-fold interquartile range; remaining participants are represented by dots. Models are arranged in the same way as in Fig. 1. Qualitative model fit, i.e., similarity to human behavior (top-right), closely mirrors quantitative model fit (Fig. 1, Table 1), with similar results for behaviors other than action bout length.

#### Most Flexible Model: “LSTM-all”

We next fit the most general ANN: a long short-term memory (LSTM; Hochreiter and Schmidhuber, 1997) model we call “LSTM-all” because it has access to “all” observable trial information *r*_*t*_ and *a*_*t*_. We trained LSTM-all to output a 4-dimensional “gist” vector *g*_*t*_(*a*) on each trial *t*, which—similar to RL_αβ_’s *v*_*t*_(*a*)—was submitted to the softmax function to obtain action probabilities (Table 1).

LSTM-all was fitted using the same CE loss as RL_αβ_ and RL_αβ_ *p f*. Hyperparameters were chosen based on visual inspection of training trajectories, and then fixed for all models.^2^ LSTM-all’s behavioral validation was outstanding, with all human behavioral markers reproduced reliably (Fig. 2B; for quantitative fit, see Fig. 1B). Hence, as expected, the constrained RL_αβ_ model does not capture human behavior as well as the flexible LSTM-all. This raises the question which specific mechanisms LSTM-all capture that RL_αβ_ was lacking.

#### Basic Value RNN: “RNN-v”

We first tested whether relaxing the functional form of the RL_αβ_ value update improves model fit. We created RNN-v, a model identical to RL_αβ_ except that it replaced *f*_*RL*_ with an unconstrained RNN *f*_*RNN*_. Notably, *f*_*RNN*_ was constrained to the same inputs as *f*_*RL*_ (i.e., did *not* recurrent states like regular RNNs; Table 1). Specifically, RNN-v implements a recurrent 3-layer MLP with 2 input units (*v*_*t*_(*a*_*t*_), *r*_*t*_), 16 hidden units (*s*_*t*_), and 1 output unit (*u*_*t*_(*a*_*t*_)). RNN-v’s free parameters θ_*RNN*_ contain weight matrices *W* and biases *b. tanh* is a non-linear activation function:

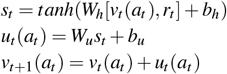

As expected, RNN-v achieved better quantitative (Fig. 1B) and qualitative fit (Fig. 2B), though the differences were relatively small.

After training, the fitted weights θ_*RNN*_ of *f*_*RNN*_ implement a value update function entirely learned from human behavior (Fig. 2A, second row). Because *f*_*RNN*_ has the same inputs and outputs as *f*_*RL*_ (Table 1), the two are directly comparable (Fig. 2A, top row). We found that when the chosen action *a*_*t*_ had a high value prior to the update (x-axis), *f*_*RNN*_ and *f*_*RL*_ were similar: Larger rewards (yellow) resulted in larger updated values than smaller rewards (blue). There was a difference with respect to reward sensitivity, however: Whereas *f*_*RL*_ showed an even spread over reward magnitudes, *f*_*RNN*_ was compressed at the high and low ends. Furthermore, when the chosen action *a*_*t*_ had a low value prior to the update, *f*_*RNN*_ diverged substantially from *f*_*RL*_: Previously low values stayed low, irrespective of the reward magnitude. In sum, allowing functional flexibility in the value update improved RNN-v’s fit to human data. The learned value update function showed a distortion in human reward sensitivity and reduced updating for low-valued actions. ^3^ However, compared to other mechanisms explored later, the improvement in RNN-v’s fit was relatively small (Fig. 1B).

#### Value RNN with Output Memory: “O-RNN-v”

We next investigated whether the ability to access not just the value of the chosen action *v*_*t*_(*a*_*t*_), but of all actions *v*_*t*_(*a*) improves model fit, creating model O-RNN-v (Table 1). Access to *v*(*a*) allows O-RNN-v to condition individual value updates on its own, as well as all other actions’ values. Indeed, model fit improved (Fig. 1B, Fig. 2B): In analyses not shown here, we observed that O-RNN-v learned to apply different value updates in different value contexts. This suggests that in the current task, humans conditioned individual value updates on the current value landscape, rather than performing them in isolation, in accordance with previous findings (e.g., Palminteri et al., 2015).

#### Value RNN with State Memory: “OS-RNN-v”

We next investigated the effects of memory more broadly, constructing OS-RNN-v, which receives its own prior hidden state *s*_*t*_ as an additional input, completing the classic RNN architecture (Table 1). Access to *s*_*t*_ allows the model to carry information between trials that is not directly related to action choice (in the way *v*(*s*) is), and can hence capture long-term contingencies or action plans (or individual differences, see Discussion). This change further improved quantitative (Fig. 1) and qualitative fit (Fig. 2B).

#### Value LSTM: “LSTM-v”

We then constructed LSTM-v, a vanilla LSTM trained on the value-based inputs (Table 1): Like the other models in the “v family”, LSTM-v submitted a 4-dimensional vector *v*(*a*) to the choice rule, and provided updates *u*_*t*_(*a*_*t*_) to the previous action (*a*_*t*_). LSTM-v is the most powerful model in the v-family, with a potentially improved capacity to capture long-term dependencies. However, model fit did not improve (Fig. 1, Fig. 2B).

#### Choice-Kernel RNNs: “-c Models”

Despite access to the full capacity of the LSTM architecture, LSTM-v does not achieve the same fit as LSTM-all (Fig. 1, Fig. 2B). The reason is that LSTM-v lacks access to the identity of the chosen action *a*_*t*_ (Table 1). We therefore investigated next how much behavioral variance in the human data could be explained by models based on action identity *a*_*t*_. To this aim, the choice-kernel or “c family” of models had access to *a*_*t*_, but *not r* (Table 1). We constructed the c family in direct correspondence to the v family, allowing direct comparison of the role of memory (differences between “O-”, “S-”, and “OS-” models).

We found that similar to value RNNs, choice-kernel RNNs benefit from access to *c* (“O” models) and *s* (“S” models”), with even larger observed differences in model fit (Fig. 1). This suggests that long-term memory might play a larger role in action-history dependent (i.e., choice kernel-based) than in reward-dependent (i.e., value-based) choice.

Interestingly, the choice kernel-based model with full memory capacity (LSTM-c) explains more variance than its value-based partner LSTM-v (Fig. 1), suggesting that in the current task, participants relied more on action patterns than reward information. This could reflect an increased use of exploratory strategies (Fintz et al., 2021) or lack of task engagement.

#### Additive Combination of Value and Choice Kernel

“▪-cv Models” We next asked how v and c processes interact. We first constructed models that calculate both choice kernels and values independently, and only (additively) combine them at the final decision stage (“additive” family, Table 1). Indeed, additive models fit the human data much better than either component model alone, at every level of memory use (Fig. 1). Additive models also show substantially better qualitative fit than prior models (Fig. 2B).

#### Interactions between Value and Choice Kernel: “-all Models”

The models in the additive family lack one crucial capability of LSTM-all, notably the non-linear combination of reward and action identity: By design, additive models cannot capture processes that combine, compare, or contextualize reward and action identity with each other because the calculations are performed separately (Table 1).

We hence constructed the final “interactive” model family to assess evidence for choice kernel-value interactions (Table 1). Interestingly, interactive models do not generally provide improvements in quantitative (Fig. 1) or qualitative fit (Fig. 2B) over additive models (with the potential exception of LSTM-all). This might suggest that humans process reward histories and action histories using distinct cognitive— and potentially neural—pathways, in accordance with previous findings (K. J. Miller et al., 2019).^4^

## Discussion

This paper addresses several open questions in the field of RL and cognitive modeling: 1) How much behaviour in typical RL tasks is explained by theoretical, hand-crafted RL models? Echoing previous findings in RL (e.g., Dezfouli, Ashtiani, et al., 2019; Dezfouli, Griffiths, et al., 2019; Fintz et al., 2021; Jaffe et al., 2022; Song et al., 2017; Yang et al., 2019) and other fields (e.g., Agrawal et al., 2020; Battleday et al., 2020; Kuperwajs et al., 2022; Peterson et al., 2021), we find that hand-crafted, cognitive RL models leave substantial amounts of behavioral variance unexplained, which can reliably be captured using more flexible, ANN-based models.

2) If hand-crafted RL does not explain human choice, which mechanisms do? We find strong evidence for an outcome-insensitive decision mechanism, which might capture exploration or habits (K. J. Miller et al., 2019). This mechanism can be inspected and interpreted based on inputoutput mappings just like the value-based mechanism (Fig. 2A), though we are not showing the results here. Surprisingly, we find no evidence for non-linear interactions between reward-based and outcome-insensitive mechanisms, suggesting separate cognitive mechanisms.

3) How do humans perform RL? RL theory specifies a particular updating rule, but other functional forms are possible. Our framework shows which form the human update takes when learned directly from data.

4) What role does memory play in human action selection? While the literature on memory and its interactions with RL is rich, it has traditionally been difficult to create hand-crafted models that encompass both processes (Collins, 2019; Daw and Shohamy, 2008). We show distinct roles of memory for reward-based and reward-insensitive choice processes.

These findings extend previous studies that have aimed to achieve interpretability when using ANNs as cognitive models. Previous suggestions range from shifting the research focus from explanation to prediction (Yarkoni and Westfall, 2017), analyzing ANNs’ hidden representations (Kriegeskorte, 2015; Schaeffer et al., 2020) or behavioral predictions (Agrawal et al., 2020; Dezfouli, Griffiths, et al., 2019; Fintz et al., 2021), directly comparing ANNs to hand-crafted cognitive models (Fintz et al., 2021), creating ANNs that expose interpretable information in their hidden state (Dezfouli, Ashtiani, et al., 2019; Dezfouli, Griffiths, et al., 2019), or integrating ANN-based representations (Battleday et al., 2020; Noti et al., 2016) or entire computational components into existing, hand-crafted models (Peterson et al., 2021). Our approach is very closely related to Peterson et al. (2021).

## Limitations and Future Work

### Dataset Size

One limitation of the current study is the dataset size. Even though at 965 participants, the dataset (Bahrami and Navajas, 2020) is an order of magnitude larger than most standard studies in the field, we observed indices that it might be too small: We calculated the test loss of our models after training on different data sizes. Our most flexible models still showed improvements leading up to the largest size (Fig. 3). Furthermore, some results were sensitive to small changes in the size of training and testing data or on participants’ assignments. We see two main solutions: collecting a larger dataset or reducing model complexity in accordance with the given dataset (e.g., by reducing the number of hidden neurons).

**Figure 3:**
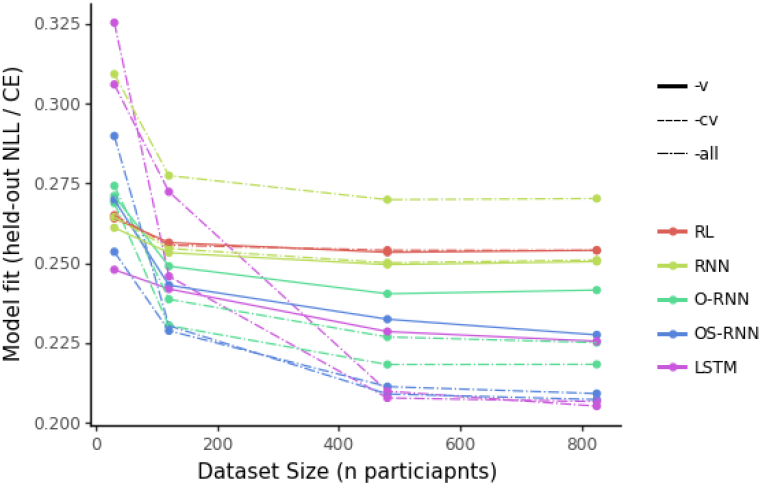
Model fits vary with training data size. Each line refers to one model, colors and linetypes specify model categories. Each model was fitted to four different datasets (30, 120, 480, or 825 participants) for 200.000 training steps, using the same hyperparameters. The trained models were then tested on the same held-out data (y-axis, negative log-likelihood / cross-entropy). More flexible models (e.g., LSTM, OS-RNN) showed greater improvements in fit than less flexible models (e.g., RL_αβ_), driven by both increased overfitting (i.e., worse fit to held-out data) when trained on smaller datasets, and superior generalization (i.e., better fit to held-out data) when trained on larger datasets.

### Task Design

Some of our results might be specific to the current task, rather than pertain to learning and decision making more broadly. For example, the strong role of choice kernel-based control might be related to the relatively large number of four alternatives, or the common closeness of alternatives to each other, which makes it difficult to identify the best (Fintz et al., 2021). If this is the case, conclusions drawn from this dataset might not explain the fundamental set-up of human learning, but rather elucidate particular task strategies (Eckstein et al., 2021; Nussenbaum and Hartley, 2019).

To alleviate this problem, future work needs to exhaustively sample the space of task features over which conclusions are aimed to be drawn (e.g., number of bandits; stochastic or deterministic reward scheme; volatile or stable trial structure; correlated, anticorrelated, uncorrelated bandits; etc.). Such a dataset would allow to marginalize over individual task features (Peterson et al., 2021), and follow a ubiquitous aspiration to introduce richer, more complex paradigms into the study of cognition (Battleday et al., 2020; Ma and Peters, 2020).

### Individual Differences

Like the majority of cognitive ANN studies, our approach does not explicitly model individual differences (however, see Dezfouli, Ashtiani, et al., 2019; Dezfouli, Griffiths, et al., 2019). This is particularly relevant in the current setting because our basic RL models have no capacity to capture individual differences, while more advanced models have this capacity as long as they are endowed with state-memory: These advanced models can adapt to individuals by conditioning decisions on their hidden state, which can represent unique individual characteristics.

## Conclusion

In this paper, we introduce “hybrid ANNs”, a combination of classic RL models and ANNs. These models have both predictive (excellent fit) and explanatory power (interpretability of the resulting models), and potentially provide detailed insight into human cognitive processes.

## Footnotes

1 We use the notation *a* to signify the vector of all actions; *a*_*t*_ refers to the action chosen at trial *t. p*(*a*) [*v*(*a*)] denotes the vector of all action probabilities [values].

2 batch size *s*_*max*_: 64; number of training steps: 200.000; size of hidden layer: 16; Adam optimizer learning rate for RL_αβ_, RL_αβ*p f*_, and all models of the “-all” family: 0.001; for all other models: 0.0001

3 Note that these patterns could conflate task-based and individual differences. For example, it is possible that some participants, e.g., due to a lack of task engagement, showed a lack of reward-informed action choice (i.e., no detectable value update in *v*_*t*+1_) and hence observed generally smaller rewards (i.e., low values *v*_*t*_). These participants might explain the left (low-value) side of the update function. A different group of participants, showing optimal task engagement, might however show great sensitivity to reward differences (i.e., easily detectable value update in *v*_*t*+1_), and hence observe consistently large rewards (i.e., high values *v*_*t*_). These participants might explain the right (high-value) side of the update function.

4 Note that this conclusion is based on the assumption that the current dataset was large enough to fit all models optimally, which might not be the case (see Discussion and Fig. 3). Some of the observed pattern could also arise if more complex models were underfit.

